# Rate of photosynthetic acclimation to fluctuating light varies widely among genotypes of wheat

**DOI:** 10.1101/435834

**Authors:** William T. Salter, Andrew M. Merchant, Richard A. Richards, Richard Trethowan, Thomas N. Buckley

## Abstract

Significant variation exists in the acclimation time of photosynthesis following dark-to-light transitions across wheat genotypes, under field and controlled conditions. Slow acclimation reduced daily carbon assimilation by up to 16%.

**Abstract:** Crop photosynthesis and yield are limited by slow photosynthetic induction in sunflecks. We quantified variation in induction kinetics across diverse genotypes of wheat for the first time. In a preliminary study using penultimate leaves of 58 genotypes grown in the field, we measured induction kinetics for maximum assimilation rate (*A*_max_) after a shift from full darkness to saturating light (1700 μmol m^−2^ s^−1^) with 1-4 replicates per genotype. We then grew 10 of these genotypes with contrasting responses in a controlled environment and quantified induction kinetics of carboxylation capacity (*V*_cmax_) from dynamic *A* vs *c*_i_ curves after a shift from low to high light (50 to 1500 μmol m^−2^ s^−1^), with 5 replicates per genotype. Within-genotype median time for 95% induction (*t*_95_) varied from 8.4 to 23.7 min across genotypes for A_max_ in field-grown penultimate leaves, and from 6.7 to 10.4 min for *V*_cmax_in chamber-grown flag leaves. Our simulations suggested that non-instantaneous acclimation reduces daily net carbon gain by up to 16%, and that breeding to speed up *V*_cmax_ induction in the slowest genotype to match that in the fastest genotype could increase daily net carbon gain by more than 4%, particularly for leaves that experience predominantly short-duration sunflecks.

## Introduction

Global food security is threatened by growing populations and diminishing increases in crop yield potential. To ensure future food security, improvements need to be made to plant yield traits that have previously been overlooked in crop breeding programs, such as dynamic properties of photosynthesis. The efficiency of photosynthetic machinery under fluctuating environmental conditions has been identified as a key target for improvement (Taylor & Long, 2017; Murchie *et al.*, 2018). In particular, the light environment of crop canopies is highly dynamic, with fluctuations occurring on the scale of seconds to minutes (Slattery *et al.*, 2018). On clear days, leaves at the top of the canopy are generally exposed to direct sunlight for the majority of the day, whilst leaves in the lower canopy rely on light in the form of sunflecks. These sunflecks can account for up to 90% of the daily available light (Pearcy, 1990).

The impact of rapid shifts in photosynthetic photon flux density (PPFD) on carbon balance is influenced by a number of physiological factors. Diffusion of CO_2_ through stomata and the mesophyll (controlled by conductances g_s_ and g_m_ respectively) can limit induction of photosynthesis (Lawson & Vialet-Chabrand, 2018), and conversely, slow deactivation of energy-consuming photoprotective mechanisms when leaves enter shadeflecks can reduce net carbon gain (Kromdijk *et al.*, 2016). Slow acclimation of photosynthesis following shade to sun transitions can also substantially reduce carbon assimilation in crop canopies. Activation of ribulose-1,5-bisphosphate carboxylase/oxygenase (Rubisco) is considered to be a critical constraint on photosynthetic induction to shade-sun transitions (Soleh *et al.*, 2016; Taylor & Long, 2017; Morales *et al.*, 2018). In a recent study, Taylor & Long (2017) predicted that carbon assimilation could be inhibited by up to 21% in wheat (Triticum aestivum) by slow induction of Rubisco in response to fluctuating light conditions. If this inefficiency could be reduced, whole canopy carbon assimilation would be improved, potentially leading to increases in yield (Long *et al.* 2006).

Screening for genetic variation in photosynthetic activation time would increase our fundamental understanding of this process and patterns in variation and dynamics of the response would allow us to identify whether this trait is a valid target for improvement through conventional breeding. However, although some variation in the kinetics of photosynthetic acclimation has been identified across soybean genotypes (Soleh *et al.*, 2016; Soleh *et al.* 2017), there is little information available regarding the diversity of this trait in wheat. Taylor and Long (2017) examined only a single genotype, partly due to the arduous nature of the “dynamic *A* vs *c*_i_” method they used to characterize the kinetics of Rubisco activation in vivo. Other less direct methods such as in vitro Rubisco assays, or high-throughput phenotyping (HTP) field techniques such as multispectral imaging, could be applied more readily to the task of phenotyping many genotypes. However, in vitro assays do not capture the interaction of diffusional and biochemical induction. Stomatal and mesophyll diffusion influence [CO_2_] in chloroplasts, which in turn regulates Rubisco activase (Portis *et al.*, 1986) – and HTP methods cannot yet quantify photosynthetic rate per se, nor its induction kinetics. Direct measurement of gas exchange in intact leaves thus remains, in our view, the only suitable method for phenotyping these traits.

We used a three-phase approach to quantify the extent of genetic variation in photosynthetic acclimation kinetics in wheat and the potential for directed breeding to enhance productivity by harnessing this variation. First, we measured the kinetics of acclimation for CO_2_-saturated photosynthesis after a shift from darkness to saturating light in penultimate leaves of 58 genotypes of field-grown wheat. Second, we then studied acclimation kinetics more intensively for a subset of 10 of these genotypes, using the dynamic *A* vs *c*_i_ method to quantify induction of RuBP carboxylation and regeneration capacities over time after a switch from low to high light. Finally, we used modeling to quantify the improvement in diurnal net carbon gain that could be achieved by breeding for faster photosynthetic induction within the observed range of variation across genotypes.

## Methods

### Plant material

Detailed analysis of photosynthetic acclimation was conducted on wheat grown under controlled conditions in June/July 2018. 10 genotypes were selected from those grown in the field the previous year (Table S1). Seed was sown in 5 l pots with compost mix containing a slow release fertiliser (Evergreen Garden Care Australia, Bella Vista, NSW, Australia). Day and night temperatures were maintained at 23.7 ± 1.7°C and 12.0 ± 1.8°C (mean ± s.d.) respectively, and relative humidity at 67.2 ± 6.1% and 74.0 ± 8.1%. Growth CO_2_ concentration was 482.2 ± 23.2 μmol mol^−1^ across the course of the experiment. Light was supplied by LED growth lamps (LX602C; Heliospectra AB, Göteborg, Sweden) and provided a PPFD of 800 μmol m^−2^ s^−1^ at the leaf surface. Seedlings were thinned to one per pot after germination. Plants were watered twice daily to field capacity. All gas exchange measurements were taken on the mid-section of a fully expanded flag leaf during heading or anthesis (the distribution of Zadoks phenological growth stages during these measurements is shown in Figure S1).

For measurement of photosynthetic acclimation in field-grown plants, wheat was planted in 2 × 6 m plots with 5 sowing rows per plot in Narrabri, NSW, Australia in late May 2017. 58 genotypes were examined here (Table S1). Measurements were made on 03-18 Sep 2017, within two weeks before or after anthesis (the distribution of Zadoks phenological stages across the field measurement campaign is shown in Figure S1).

### Detailed analysis of acclimation to light in flag leaves of wheat grown in a controlled environment

Plants were moved from the controlled environment room to a temperature stable laboratory at 25°C. Photosynthetic light response curves were recorded using a LI-6400 gas exchange system (Li-Cor, Lincoln, NE, USA) on one plant of each genotype. Leaves were equilibrated to chamber conditions (leaf temperature 25°C; leaf vapour pressure deficit 1.0-1.5 kPa; cuvette CO (*c*_a_) 400 μmol mol^−1^; and PPFD 1500 μmol m^−2^ s^−1^ provided by LEDs in the chamber head) for at least 40 min to allow them to reach steady state. PPFD was then reduced through 1200, 1000, 800, 600, 500, 400, 300, 200, 150, 100, 50 and 0 μmol m^−2^ s^−1^, with measurements taken immediately after chamber conditions had stabilized at each level. Light responses curves were fitted to a non-rectangular hyperbola model using nonlinear least squares in R (nls; R Language and Environment); i.e., the lesser root *A* of

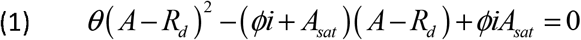

where *i* is PPFD, *A*_sat_ is the asymptotic limit of A at high PPFD, θ is a dimensionless parameter < 1, *ϕ* is the initial slope of *A*_*eq*_ vs *i*, and *R*_*d*_ is day respiration rate. *A*_*sat*_, *θ*,*ϕ* and *R*_*d*_ were fitted empirically.

Dynamic *A* vs *c*_i_ responses were recorded using four Walz GFS-3000 gas exchange systems (Heinz Walz GmbH, Effeltrich, Germany), using the method of Taylor & Long (2017). The acclimation of photosynthesis following transition from shade to saturating light was measured at a number of different *c*_a_ values, and composite *A vs c*_i_ curves were generated for each relative time point during induction. Leaf temperature was held at 25°C and VPD_leaf_ at 1.0 kPa. Each leaf was first brought to a steady state at *c*_a_ 400 μmol mol^−1^ and PPFD 1500 μmol m^−2^ s^−1^ (found to be saturating in our light response curves) over 40 min, and PPFD in the leaf chamber was then dropped to 50 μmol m^−2^ s^−1^ for 30 min. During this ‘dark phase’ the *c*_a_ was also reduced to 100 μmol mol^−1^ to inhibit stomatal closure, as per guidelines in Taylor & Long (2017). Prior to induction the *c*_a_ was increased to the desired value for induction. Induction of photosynthesis was initiated with a step change to PPFD 1500 μmol m^−2^ s^−1^, and measurements recorded every 10 seconds for 15 min. This 30 min dark – 15 min light cycle was repeated at induction *c*_a_ of 50, 100, 200, 300, 400, 500, 600, 800 and 1000 μmol mol^−1^.

*A* vs *c*_i_ curves were generated for each 10 s time point after induction of photosynthesis. The Farquhar *et al.* (1980) photosynthesis model was fitted to these curves using the ‘plantecophys’ package in R (bilinear fitting method; Duursma, 2015) to provide estimates of Rubisco carboxylation capacity (*V*_cmax_) and electron transport rate (*J*). *R*_*d*_ was set at 1.9 μmol m^−2^ s^−1^ and θ was set at 0.67 (from our light response curves). We also provided the Michaelis-Menten coefficient and the photorespiratory compensation point, calculated from the mean leaf temperature as per Bernacchi *et al.* (2001). Temperature corrections were performed during the fitting process to provide values of *V*_cmax_ and *J* at 25°C. The script used for curve fitting is provided in Supporting Information File S2.

For each leaf, we modeled the observed timecourse of induction of *V*_cmax_ using a two-phase exponential function of time:

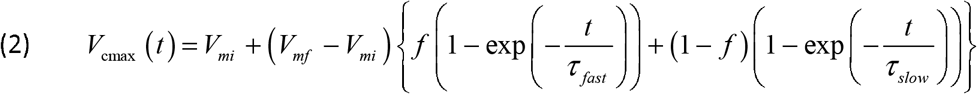

where the parameters (*V*_*mi*_ and *V*_*mf*_ are the initial and final (fully acclimated) values of *V*_cmax_, *τ*_fast_ and *τ*_slow_ slow are time constants for fast and slow phases of acclimation, respectively, and *f* is a weighting factor between zero and one) were estimated by using Solver in MS Excel to minimize the sum of squared differences between measured and modeled *V*_cmax_. We found that this two-phase model produced a substantially better fit to our data than a single-phase model as used by Taylor and Long (2017), with *r*^2^ ranging from 0.957 to 0.996 (median = 0.990).

### Photosynthetic acclimation to light in penultimate leaves of field grown wheat

We used OCTOflux to measure photosynthetic acclimation upon transition from darkness to saturating light in penultimate leaves of 58 genotypes of field grown wheat. This system is described elsewhere (Salter *et al.*, 2018). It is an open-flow single-pass differential gas exchange system with eight leaf chambers (5 × 11 cm), designed to maximize throughput for measurements of CO_2_-and light-saturated net photosynthesis rate (*A*_max_). Each chamber has a white LED light source above the adaxial leaf surface, a Propafilm window, four small mixing fans and a type T thermocouple kept appressed to the abaxial surface. Stable dry air is created by mixing CO_2_ and dry air from pressurized cylinders with mass flow controllers into a buffering volume (~40 L) containing a powerful fan. This gas is then split into nine streams: a reference stream, which flows through the reference cell of a differential infrared gas analyzer (IRGA; Li-7000, Li-Cor, Lincoln, Nebraska), and eight sample streams, each of which runs through a mass flow meter to a leaf chamber and back to the IRGA, where it is either vented to the atmosphere or directed through the IRGA sample cell, using solenoid valves.

Tillers were cut in the field, immediately recut under distilled water and placed into darkness and transported by vehicle to the laboratory (about 1 km away; time from cutting to laboratory was 5-15 min), and kept in darkness for a further 0 – 30 minutes before measurement. Each leaf was enclosed in a leaf chamber and exposed to saturating PPFD (1700 μmol m^−2^ s^−1^) and ambient CO_2_ (4800 – 5000 μmol mol^−1^), and then allowed to acclimate to these conditions. To verify that *A*_max_ measured at these high *c*_a_ values did not differ substantially from the true *A*_max_, which occurs at the transition point between RuBP-regeneration-limited and triose phosphate utilization (TPU) limited photosynthesis, we measured traditional *A* vs *c*_i_ curves in 18 leaves and extrapolated these to high *c*_a_ using a biochemical model (Farquhar *et al.* 1980) as extended by Busch *et al.* (2018), and found that A_max_ at 5000 μmol mol^−1^ was an excellent proxy for true *A*_max_ (r^2^ = 0.9841, slope = 0.9968; Figure S2). Full details of these tests are given in Salter *et al.* (2018).

The present study took advantage of the fact that the sample gas stream from one of the eight chambers could be continuously measured during photosynthetic acclimation to saturating light. We recorded net CO_2_ assimilation rate every two seconds until stability was achieved (average ~14 min), and the record of A vs time was then modeled with the following sigmoidal equation:

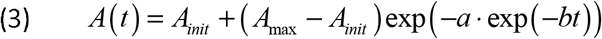

where *A*_*init*_, *A*_max_, *a* and *b* are positive empirical parameters fitted by using Solver (GRG nonlinear engine) in Microsoft Excel to minimize the sum of squared differences between measured and modeled A. The times for A to rise by 25%, 75% and 95% of the difference between *A*_*init*_ and *A*_max_ (*t*_25_, *t*_75_ and *t*_95_, respectively) were then calculated from the fitted parameters, as *t*_*x*_ = ln(*a*/ln(1/[0.01·*x*]))/*b*, where *x* = 25, 75 or 95. The “rise time,” or the time required for A to increase through the middle 50% of its dynamic range, was calculated as *t*_75_ – *t*_25_.

Because the workflow was organized around the broader phenotyping study (in which 160 genotypes were measured), replication was unbalanced among the 58 genotypes for which we recorded acclimation kinetics, with *n* = 1 to 4 replicate plants per genotype.

### Modeling impact of acclimation kinetics on carbon gain

We simulated the impact of observed variability in photosynthetic acclimation kinetics on diurnal carbon gain for different sunfleck lengths and canopy positions, using a modeling approach similar to that of Taylor and Long (2017), and simulating *V*_cmax_ induction kinetics using median kinetic parameters for the slowest and fastest of the 10 genotypes studied. To assess the role of sunfleck length and canopy position, we calculated irradiance based on expressions given by Retkute *et al.* (2018) and de Pury and Farquhar (de Pury & Farquhar, 1997), rather than the sample ray tracing model output used by Taylor and Long (2017). The modeling approach is described in detail in Appendix S1; sample diurnal traces of leaf irradiance, including alternating sunflecks and shadeflecks, are shown in Figure 1.

**Figure 1.**
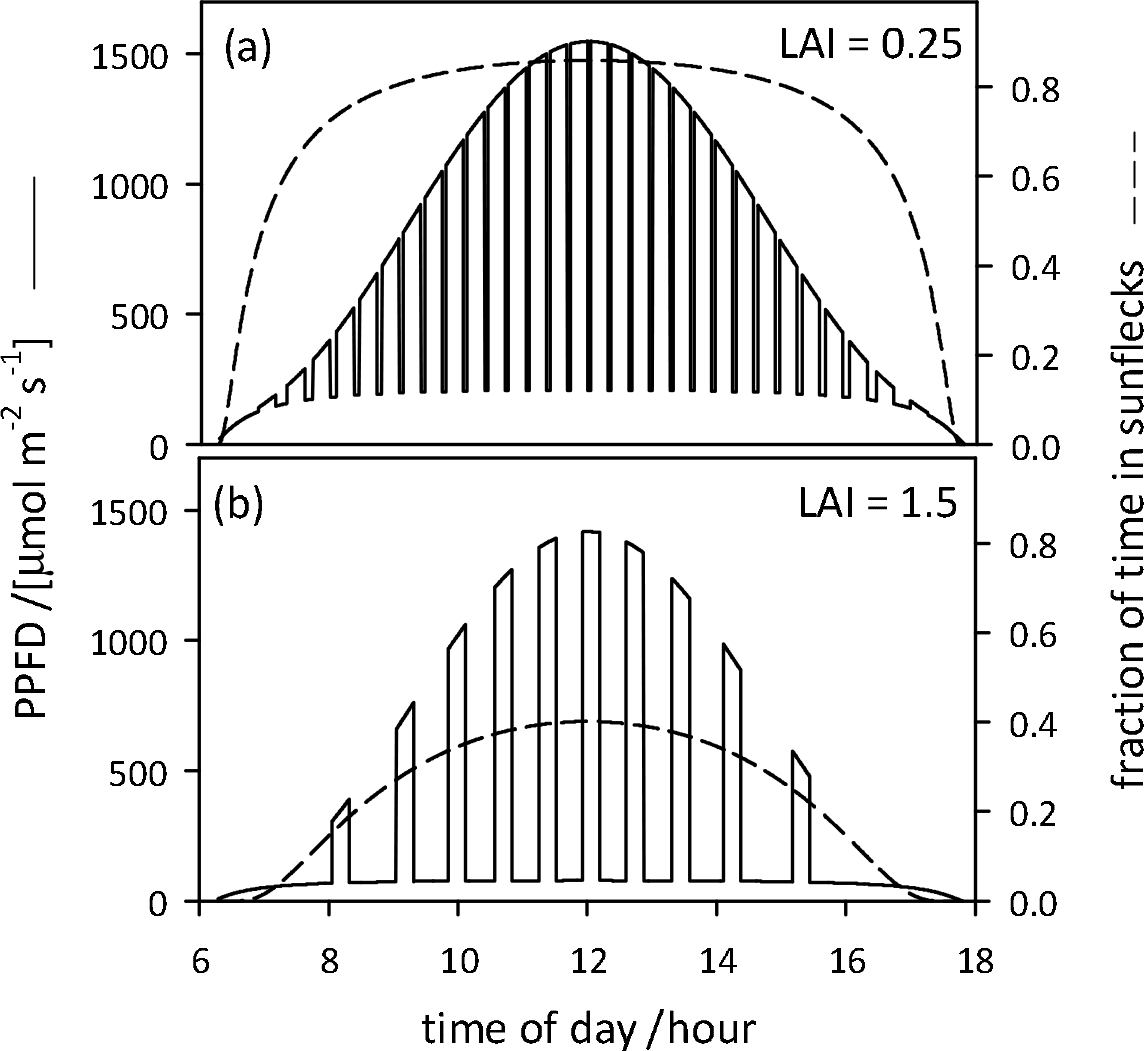
Representative time-courses of simulated PPFD with alternating sunflecks and shadeflecks (solid lines, left axis) and the fraction of time spent in sunflecks by leaves (dashed lines, right axis), for two canopy positions: cumulative leaf area indices of (a) 0.25 m^2^ m^−2^, and (b) 1.5 m^2^ m^−2^. Simulations assumed a constant sunfleck duration of 16 minutes.

### Analysis of *A* vs *c*_i_ data with the ‘one-point method’

We tested whether the ‘one-point method’ of De Kauwe *et al.* (2016) would provide robust estimates of *V*_cmax_ during photosynthetic acclimation to dark-to-light transitions. For this analysis we used the dark-to-light photosynthetic induction at c_a_ 400 μmol mol^−1^ from our dynamic *A* vs *c*_i_ analysis, and estimated *V*_cmax_ by fitting the Farquhar *et al.* (1980) model to the data using the ‘plantecophys’ package in R (Duursma, 2015).

### Statistical analysis

We tested for differences among genotypes in functional parameters of acclimation kinetics (*t*_95_ and *t*_75_-*t*_25_) using analysis of variance (function aov() in base R) with genotype as a categorical independent variable and *t*_95_, etc., as dependent variables (after transformation to improve normality, by inversion [i.e., *y* = 1/*t*_95_] for chamber-grown plants, and by log transformation for field-grown plants). Outliers for *t*_95_ and *t*_75_-*t*_25_ were removed from the *V*_cmax_ dataset on the basis of a Grubbs test (R package ‘outliers’) applied to each genotype; this resulted in removal of three values for *t*_95_ and four for *t*_75_-*t*_25_.

## Results

### Measurement and modeling of acclimation kinetics

A vs c_i_ curves fit to each 10 s interval following transition from shade (50 μmol m^−2^ s^−1^) to saturating light (1500 μmol m^−2^ s^−1^) revealed limitations imposed by *V*_cmax_ at lower *c*_i_ and by *J* at higher *c*_i_ throughout induction. Whilst specific acclimation times varied between individual leaves and among genotypes (fitted dynamic *A* vs *c*_i_ curves for a slow and a fast acclimating leaf are shown in Fig 2a and 2b respectively), general trends in induction kinetics were clear (Fig S3). Both *V*_cmax_ and *J* increased immediately after transition to saturating light, however *J* increased more rapidly than *V*_cmax_ in the first three minutes and also saturated more quickly. As a result, *c*_i,trans_ (the *c*_i_ at which the primary limitation imposed on photosynthesis switches between *V*_cmax_ and *J*) rose to a maximum of 563 ± 3.2 μmol mol^−1^ three minutes after transition to saturating light, decreasing to 422 ± 4.1 μmol mol^−1^ after seven minutes and remaining relatively stable after this time (Fig S3c). The high values of *c*_i,trans_ throughout induction indicate that *V*_cmax_ is likely always limiting to photosynthesis under field conditions (assuming *c*_a_ of approx. 400 μmol mol^−1^). A two-phase exponential model was fitted to the measured *V*_cmax_ data with *r*^2^ ranging from 0.957 to 0.996 (median = 0.990), a representative timecourse of *V*_cmax_ induction is shown in Figure 3b.

**Figure 2.**
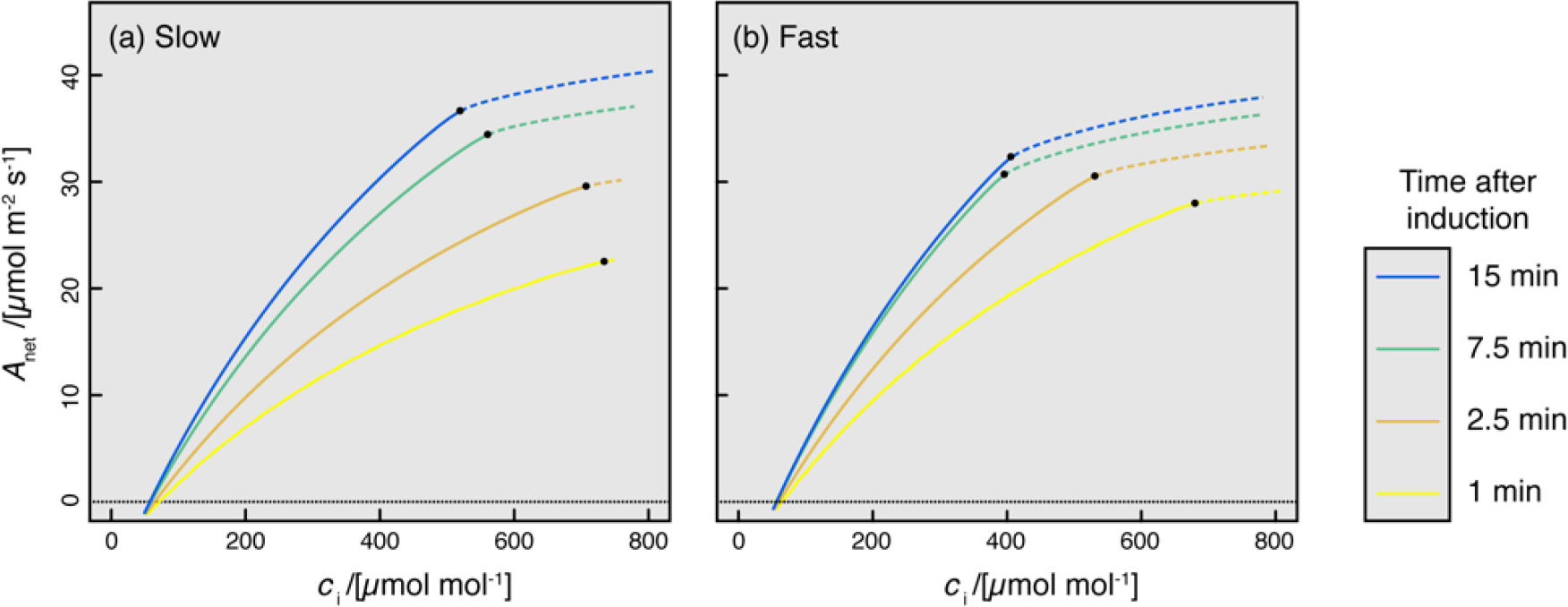
Two examples of dynamic *A* vs *c*_i_ curves, for a leaf with relatively slow acclimation of photosynthesis to light (a; time for *V*_cmax_ to increase by 95% of the difference from its initial value to its final value, *t*_95_ = 12.5 min), and a leaf with faster acclimation (b; *t*_95_ = 5.8 min). In each panel, each curve comprises a Rubisco carboxylation-limited segment (solid lines) and an RuBP regeneration-limited segment (dashed lines), and four curves are shown, each corresponding to a different time after exposure to saturating PPFD (yellow: 1 min; orange: 2.5 min; green: 7.5 min; blue: 15 min).

**Figure 3.**
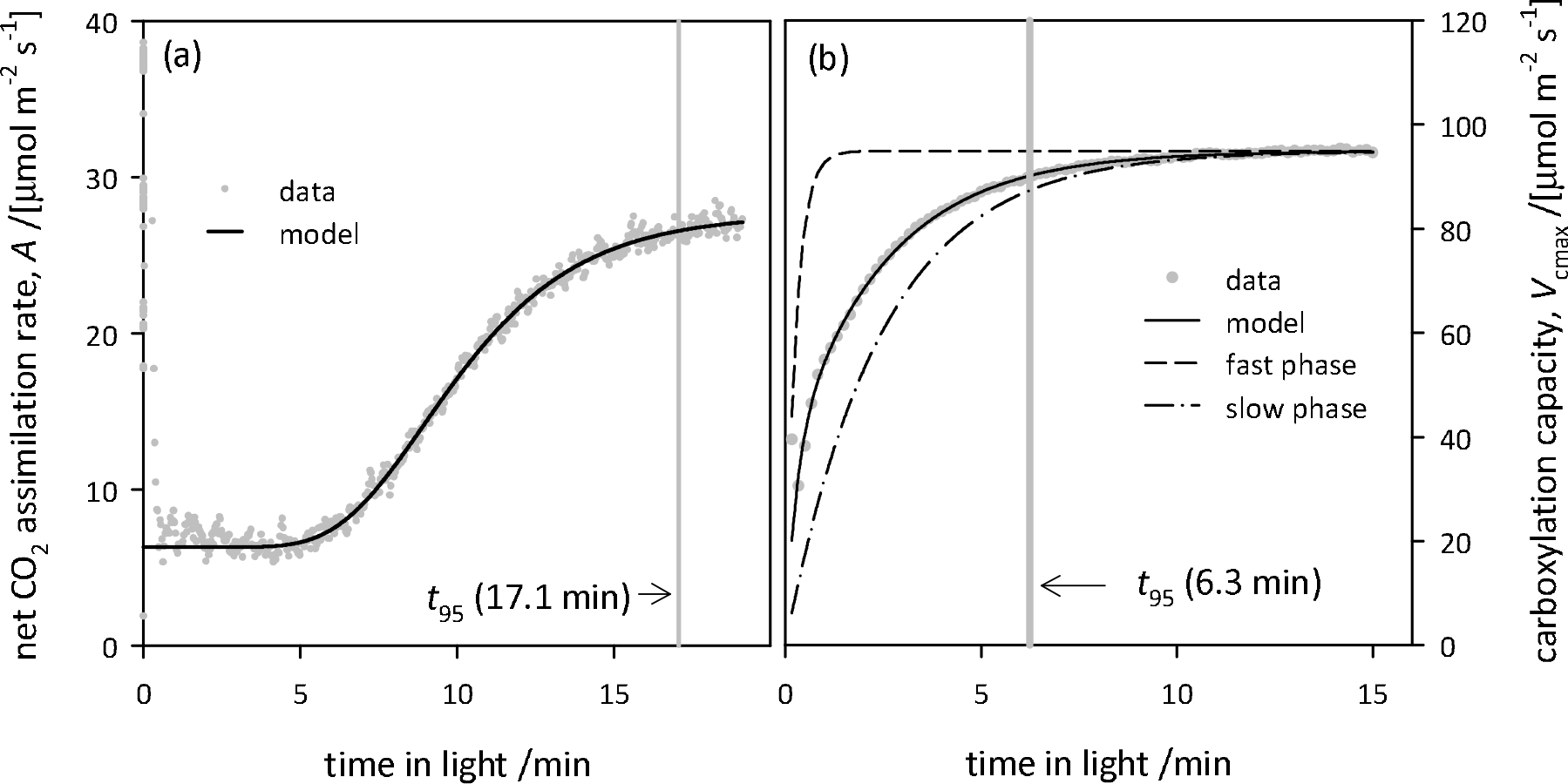
Representative time-courses of CO_2_-and light-saturated net assimilation rate measured in field-grown plants (*A*_max_; a) and carboxylation capacity inferred from dynamic *A* vs *c*_i_ curves measured on chamber-grown plants (*V*_cmax_; b). The time at which *A*_max_ or *V*_cmax_ rose through 95% of its dynamic range (*t*_95_) is shown with a vertical grey bar in both panels. Solid black lines indicate model fits (a: Eqn 3; b: Eqn 2); in (b), the dashed and dash-dot lines represent the fast and slow phases of the model for *V*_cmax_ induction, respectively (each adjusted to the same asymptote as the full model).

Photosynthetic induction kinetics of plants grown under field conditions differed from those grown under controlled conditions, specifically there was an initial lag phase after transition to saturating light. A representative timecourse of *A*_max_ induction of field grown plants is shown in Figure 3a. A sigmoidal equation was fitted to the acclimation kinetics of *A*_max_ of field grown plants with median *r*^2^ > 0.99.

### Variation in photosynthetic induction kinetics

For *A*_max_ in penultimate leaves of field-grown wheat, within-genotype median *t*_95_ (the time for *A*_max_ to rise through 95% of its dynamic range) ranged from 8.4 to 23.7 min across genotypes (Fig 4a). The within-genotype median for *t*_75_ – *t*_25_ (the time required for *A*_max_ to increase through the middle 50% of its dynamic range) varied from 1.5 to 7.6 min (Fig 4b). Differences among genotypes were not significant for either variable (F(57,73) = 0.8, p = 0.81 for *t*_95_, and F(57,73) = 0.94, p = 0.6 for *t*_75_ – *t*_25_). Across genotypes the final *A*_max_ was unrelated to *t*_95_ (*r*^2^ = 0.027, p = 0.058) or *t*_75_-*t*_25_ (*r*^2^ < 0.001, p =0.923) (Fig S4a and S4c).

The rate of induction also varied greatly across genotypes for *V*_cmax_ in flag leaves of chamber-grown wheat. Within-genotype medians ranged from 6.7 to 10.4 for *t*_95_, and from 2.2 to 3.1 for *t*_75_ – *t*_25_ (Fig 4c,d). Differences among genotypes were highly significant for *t*_95_ (*F*(9,37) = 3.97, *p* = 0.0013), but not significant for *t*_75_ – *t*_25_ (*F*(9,37) = 1.99, *p* = 0.07). The corresponding within-genotype median time constants for the fast and slow phases of acclimation, *τ*_fast_ and *τ*_slow_, respectively, ranged from 0.05 min to 0.51 min (*τ*_fast_) and from 2.6 to 4.5 min (*τ*_slow_); the weighting factor for the fast phase (*f*) was 0.39 in the fastest genotype and 0.49 in the slowest. Figure 5 shows representative time-courses of acclimation of normalized *V*_cmax_ corresponding to these lower and upper deciles for *τ*_fast_ and *τ*_slow_. The final *V*_cmax_ was unrelated to *t*_95_ (*r*^2^ = 0.072, *p* = 0.060) but was loosely correlated with *t*_75_ – *t*_25_ (*r*^2^ = 0.146, *p* = 0.006) (Fig S4b and S4d).

**Figure 4.**
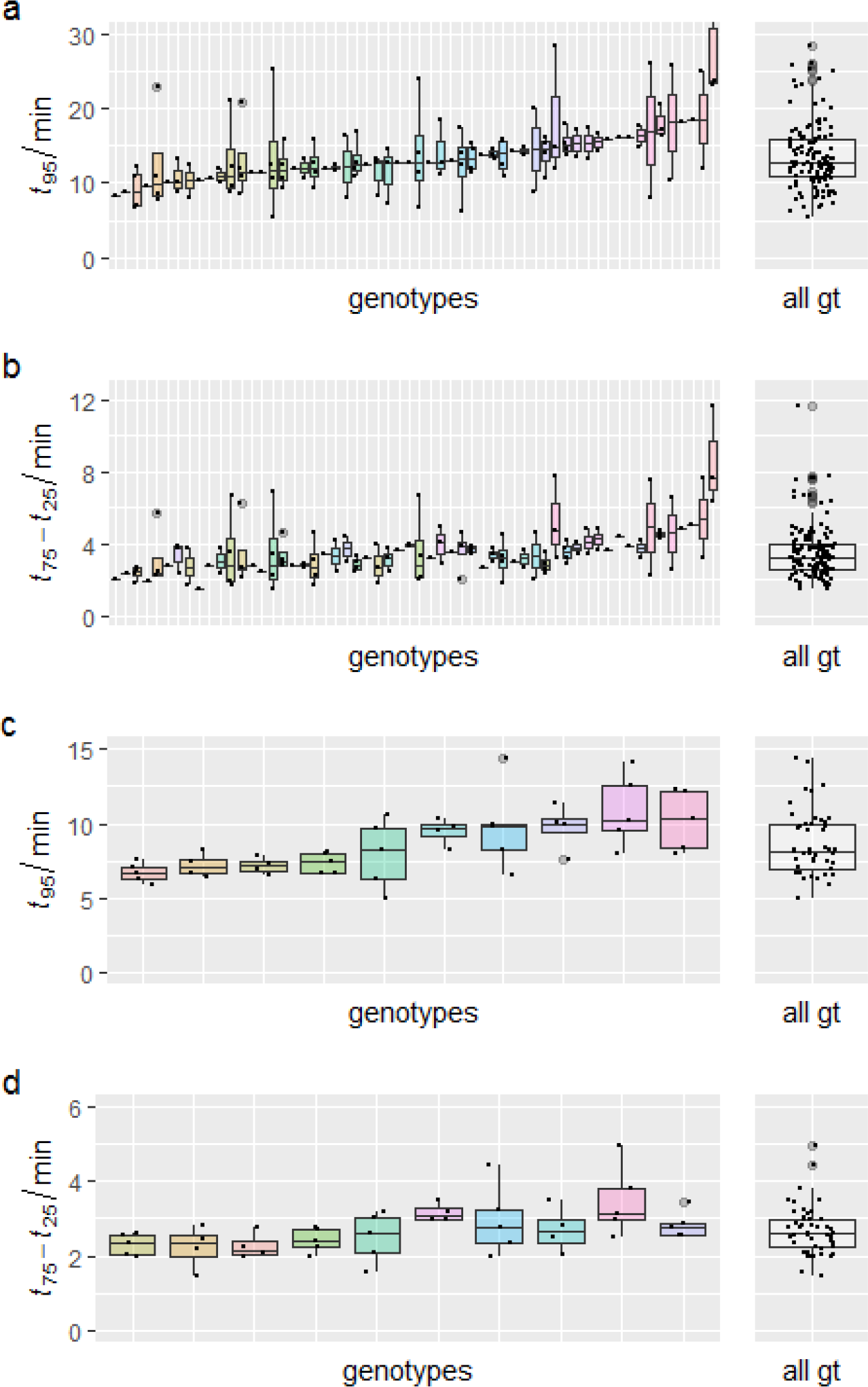
Distribution of values of (a,c) *t*_95_ and (b,d) *t*_75_ - *t*_25_, the time for Amax (a,b) or Vcmax (c,d) to increase through 95% of its dynamic range (*t*_95_), or through the middle 50% of its dynamic range (*t*_75_ - *t*_25_). 58 genotypes were studied for (a) and (b), of which 37 had 2-4 replicates; 10 genotypes were studied for (c) and (d), each with 5 replicates. The center line in each box plot indicates the median, the upper and lower bounds of each box indicate the 75th and 25th percentiles, respectively, the whiskers indicate the 90th and 10th percentiles, respectively, and the black circles indicate individual values above or below the latter percentiles. Distributions for all genotypes combined are shown at right.

**Figure 5.**
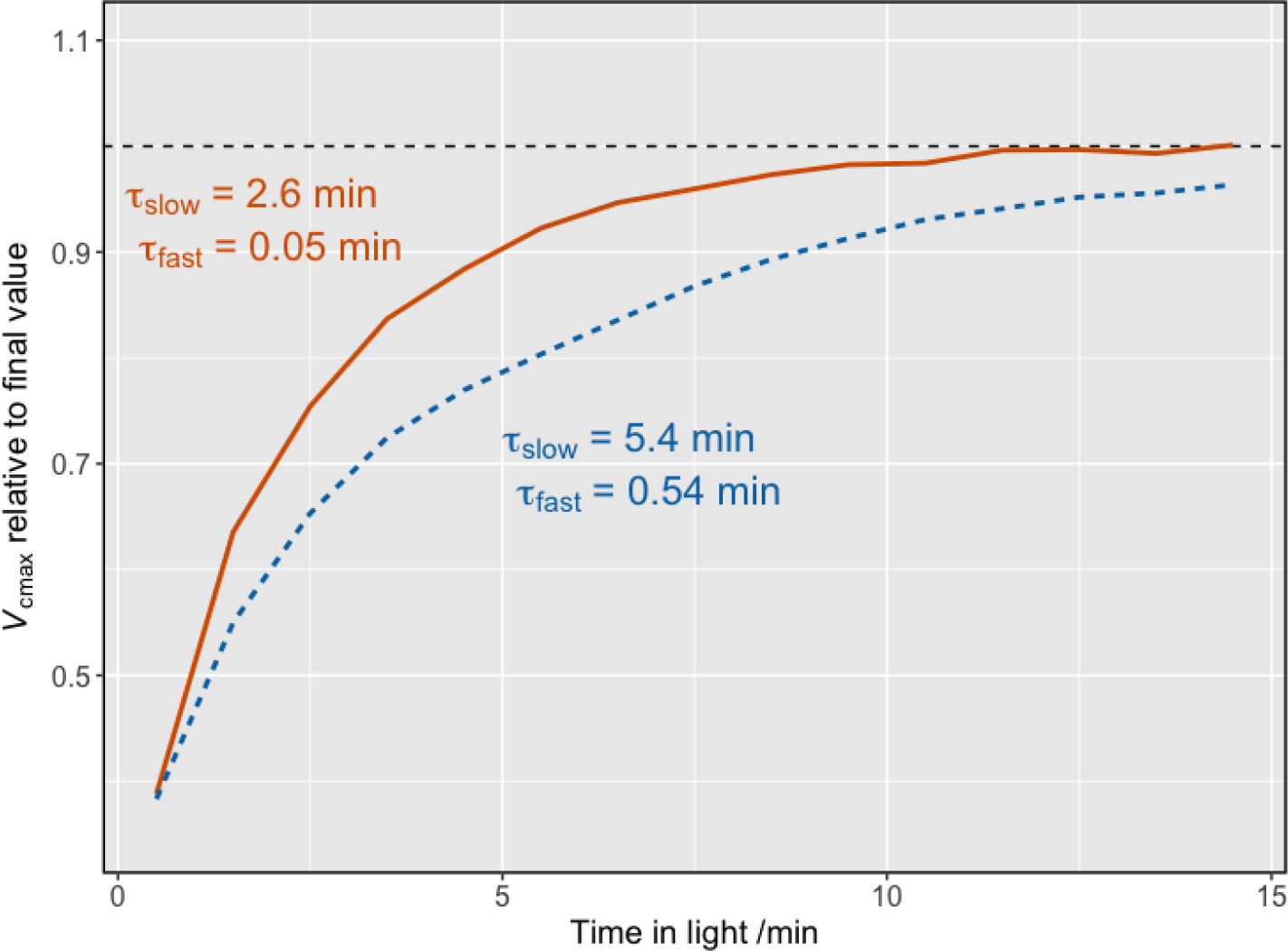
Time-courses for acclimation of normalized *V*_cmax_ for leaves using the lower decile (orange line) and upper decile (blue dashed line) for the time constants, *τ*_slow_ and *τ*_fast_, of the slow and fast phases of *V*_cmax_ acclimation. Blue line: *τ*_slow_ = 5.4 min, *τ*_fast_ = 0.54 min; orange line: *τ*_slow_ = 2.6 min, *τ*_fast_ = 0.05 min.

### Simulated effect of variation in acclimation kinetics on diurnal carbon gain

Our simulations predicted that non-instantaneous acclimation of photosynthesis to sunflecks could reduce daily carbon gain by as much as 16% (Figure 6a). The reduction was generally greatest for shorter-duration sunflecks, because photosynthesis has less opportunity to approach its fully acclimated “target” value during short sunflecks. This reduction was greater for the slowest genotype than for the fastest under most conditions, with the exception of short-duration sunflecks in upper-canopy leaves (LAI = 0.25 m^2^ m^−2^) (Figure 6a). Leaf orientation had fairly small effects on simulated carbon losses due to slow induction (Figure S5), so results presented in the main text were integrated over a spherical leaf angle distribution.

**Figure 6.**
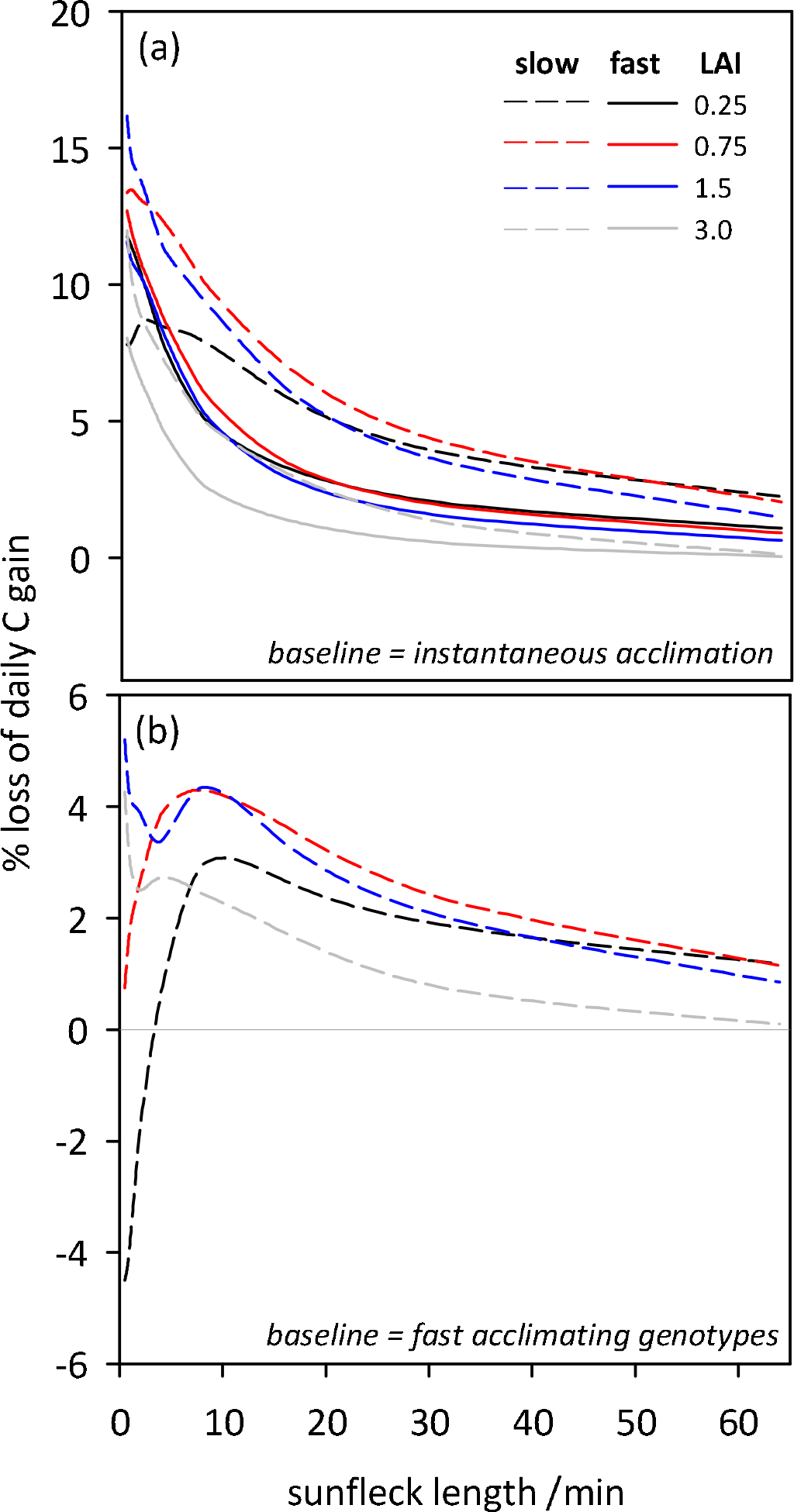
Simulated percent loss of potential total diurnal carbon gain caused by slow acclimation of photosynthesis to fluctuating light. In (a), the baseline for comparison is instantaneous acclimation, and the solid and dashed lines are results using the 10th and 90th percentiles, respectively, of kinetic parameters (*τ*_slow_ and *τ*_fast_, the time constants for the slow and fast phases of acclimation of *V*_cmax_) to model acclimation. In (b), the baseline for comparison is the faster-acclimating genotypes represented by solid lines in (a). Black, red, blue and grey lines are simulations at four different canopy depths (0.25, 0.75, 1.5 and 3.0 m^2^ m^−2^ cumulative leaf area index, respectively). All results are averaged across a spherical leaf angle distribution using Monte Carlo sampling, as described in the main text.

To consider the gains that could realistically be achieved by breeding, given the variability we observed, we also computed the % loss of daily carbon gain using the “fast” acclimating genotypes as the baseline for comparison with “slow” genotypes, rather than using instantaneous acclimation as the baseline (Fig 6b). The relative advantage of faster acclimation was greatest for sunflecks of short to intermediate duration; for example, for 8-min sunflecks, “slow” genotypes gained 2.9 to 4.3% less carbon over a day than “fast” genotypes.

## Discussion

We found greater than two-fold variation across genotypes of wheat in the time required for 95% photosynthetic induction (*t*_95_) after exposure to saturating light, both for maximum photosynthesis rate in penultimate leaves and more specifically for carboxylation capacity in flag leaves. Our simulations suggest that diurnal carbon gain is depressed by up to 15% by non-instantaneous induction of photosynthesis in sunflecks, and is up to 4% lower in the “slowest” genotypes that we studied as compared to the “fastest”. This complements recent work (Taylor & Long, 2017) documenting the potential impacts on carbon gain of slow Rubisco induction in sunflecks by demonstrating variation in this important trait in available genetic resources, showing that realistic gains are achievable even using traditional breeding.

### Variation in induction kinetics

Our preliminary analysis of 58 wheat genotypes suggested wide variation in acclimation kinetics, with genotype median *t*_95_ varying 2.82-fold overall (2.66-fold among the 37 genotypes for which we had at least two measurements), and our laboratory study with balanced replication (n=5) within ten genotypes found 1.6-fold variation in genotype median *t*_95_ for *V*_cmax_ induction. Up till now there has been little information about diversity of photosynthetic induction kinetics across wheat genotypes, however Soleh *et al.* (2017) found wide variation across 37 soybean cultivars and noted that this variation was genetically determined (i.e. stable across different leaf positions and phenological stages). As in our study in wheat, differences among soybean genotypes were attributed largely to variation in the rate of Rubisco activation. Additionally, induction kinetics were not correlated with steady-state photosynthetic capacity, and there was little evidence for this in our study (Fig S4). This observation, if held true over a broader range of studies has large ramifications for breeding approaches based upon the magnitude of *A*_max_, in particular those focused solely on the flag leaf. Based on evidence presented here, efforts to improve net carbon capture across canopies must also consider the responses of A to short term changes in the environment as a dynamic acclimation property that is at least partially genetically determined (Murchie *et al.* 2018).

Acclimation of *A*_max_ in field-grown penultimate leaves differed markedly from acclimation of *V*_cmax_ in chamber-grown flag leaves. The former was sigmoidal with time, having a lag phase and longer *t*_95_ (median 12.62 min), whereas the latter was double-exponential with time, with no lag and shorter *t*_95_ (median 8.25 min). These differences are not necessarily surprising, given the numerous differences between the two experiments; we initially discovered wide variation in *t*_95_ for *A*_max_ in the field experiment, and then designed the *V*_cmax_ experiment to be comparable to that of Taylor and Long (2017) rather than the *A*_max_ experiment. We note, however, that the ‘rise time’ (*t*_75_ – *t*_25_, the time required for *A*_max_ or *V*_cmax_ to rise through the middle 50% of its dynamic range) was more similar between the two experiments (median 3.26 min for *A*_max_ vs 2.60 min for *V*_cmax_) than was *t*_95_, which may suggest that different processes give rise to the lag and rise phases. Future work should test for effects of leaf rank and growth environment under otherwise identical conditions, and should aim to determine whether the lag phase observed in *A*_max_ induction occurs when leaves are pre-acclimated in low PPFD rather than darkness, as is more realistic for leaves in a natural canopy.

Photosynthetic induction has long been known to involve at least two phases. The initial, fast phase to involve availability of RuBP or other Calvin cycle intermediates and is complete within 1-2 minutes (Pearcy 1990), which is consistent with the median *t*_95_ that we found for the fast phase of *V*_cmax_ induction (within-genotype median *τ*_fast_ = 19.1 sec, which gives *t*_95_ = 57 sec). The slower phase apparently involves light-dependent activation of Rubisco by Rubisco activase (Rca), with time constants of 4-5 min reported for Alocasia and Spinacia oleracea (Pearcy 1990) (cf. 2.6 – 5.6 min for *τ*_slow_ in this study). In low light, sugar phosphates bind to Rubisco active sites, inhibiting carboxylation of RuBP. To restore normal function, Rubisco activase (Rca) uses energy from ATP hydrolysis to actively remove these inhibitors. Rca is sensitive to the chloroplast ADP/ATP ratio and redox status and so mediates Rubisco activation in response to light (Carmo-Silva *et al.* 2015). The variation in Rubisco activation kinetics found among wheat genotypes in our study could possibly be attributed to differences in (1) the total concentration of Rca, (2) the relative concentrations of the α-and β-Rca isoforms to each other, (3) the binding affinities of Rubisco to inhibitors and of Rca to Rubisco, and (4) the localization of Rca relative to Rubisco. In rice, Rca overexpressing mutants maintain higher Rubisco activation states in the dark and respond more quickly to changes in the light environment than wild type plants (Yamori *et al.* 2012). Arabidopsis mutants expressing only the β-Rca isoform – less sensitive to chloroplast redox status and ADP/ATP ratio than α-Rca – had faster photosynthetic induction rates and exhibited increased growth under fluctuating light compared to plants with both isoforms (Carmo-Silva & Salvucci 2013). As for the binding affinities and co-localization of these two enzymes, much less is known (for a review of current knowledge please see Mueller-Cajar *et al.* 2014) and future work should seek to address these gaps in knowledge. The recent characterization of the wheat Rca gene structure, as well as advances in genomic, proteomic and transcriptomic techniques, should provide a better understanding of these limitations and allow for a more targeted breeding approach to improve photosynthesis under dynamic light conditions (Carmo-Silva *et al.* 2015).

We did not measure deacclimation kinetics, but for modeling we assumed deacclimation kinetics in shadeflecks to be slower than acclimation in sunflecks by a factor of 5/3 (1.67), following Taylor and Long (2017). However, other evidence suggests deacclimation kinetics may be much slower still (e.g., 22-30 min in Alocasia and Spinacia; Pearcy 1990), which may mitigate the inferred benefits of breeding for faster acclimation, as discussed below.

### Impact of slow induction on photosynthesis

Consistent with the recent report by Taylor and Long (2017), our modeling found that non-instantaneous acclimation of photosynthesis to fluctuating light can reduce daily carbon gain by up to 15%. The present study extends that conclusion by quantifying the potential impact of varying sunfleck duration, canopy position (which influences the relative proportions of time spent by leaves in sunflecks vs. shadeflecks) and leaf orientation. Specifically, we found that the impact of slow acclimation was greatest for short sunflecks, because when sunflecks are similar to or much longer than the *t*_95_ for acclimation, leaves will be fully acclimated for most of each sunfleck, and thus losing little potential carbon gain. Leaf orientation had very small effects (Fig S5). The effect of canopy position was also fairly small in most cases (Fig 6), though projected % carbon losses due to slow acclimation were generally greatest at intermediate canopy depths (cumulative LAI = 0.75 m^2^ m^−2^).

An important exception was for leaves of slow-acclimating genotypes in upper canopy layers (LAI = 0.25 m^2^ m^−2^), which our simulations suggested would experience only half as much daily carbon loss as lower-canopy leaves (LAI = 1.5 m^2^ m^−2^) (7.6 vs 15.9%) for very short sunflecks (< 1 min). In fact, for upper-canopy leaves with very short sunflecks, slow-acclimating genotypes actually had a slight advantage over fast genotypes in our simulations (dashed black line in Fig 6b). This is a consequence of the wider range that we observed (and which we used to drive the simulations) for *τ*_fast_ (upper and lower deciles = 0.54 and 0.05 min, respectively) vs *τ*_slow_ (5.6 vs 2.6 min). This meant that the faster and thus greater downregulation in “fast” genotypes during short shadeflecks left them at a lower initial photosynthetic rate at the start of the subsequent sunfleck than “slow” genotypes, which outweighed the benefits of their faster upregulation during sunflecks (Fig 7a). For longer shadeflecks, the reverse was true (Fig 7b). However, we emphasize that this result depends heavily on the assumed kinetics of photosynthetic deacclimation during shadeflecks; as noted earlier, we assumed, following Taylor and Long (2017), that the time constants for deacclimation were 1.67 times greater than for acclimation. Had we instead assumed deacclimation to be 4-6 times slower than acclimation, as suggested by some data (Seeman et al 1988; Woodrow & Mott 1989; Way & Pearcy 2012), it is unlikely that 'slow' acclimating genotypes would have a carbon gain advantage except perhaps in extremely brief sunflecks. This highlights the need for future work to quantify the kinetics of deacclimation.

**Figure 7.**
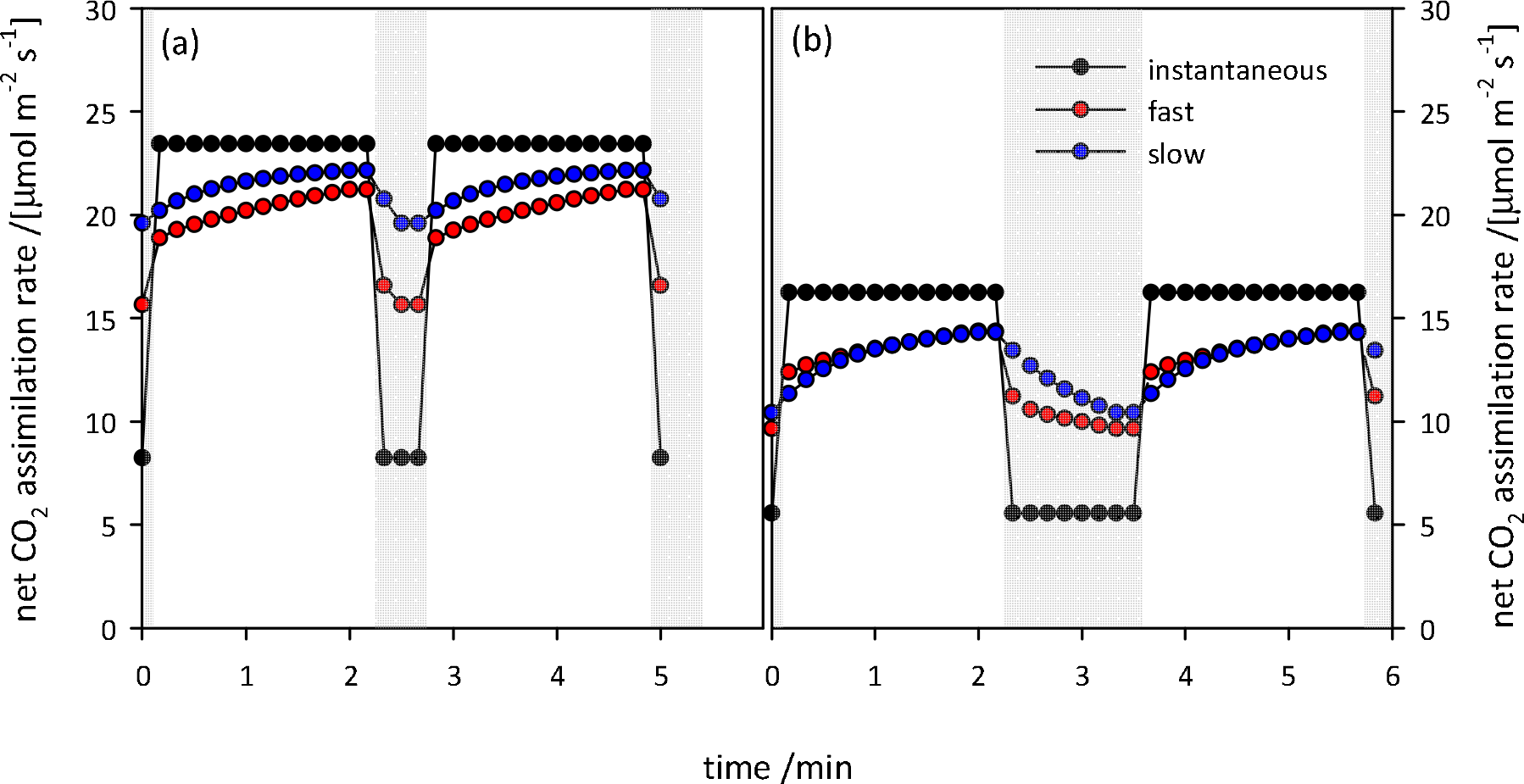
Simulated dynamics of net assimilation rate during sunflecks, and the acclimated target value of assimilation rate during shadeflecks, showing that fast-acclimating leaves (red symbols) are actually at a disadvantage relative to slow-acclimating leaves (blue symbols) when sunflecks are long and shadeflecks are short, as in panel (a). Simulations are show for horizontal leaves at mid-day at two canopy positions: cumulative leaf area indices of (a) 0.25 m^2^ m^−2^, and (b) 1.5 m^2^ m^−2^. Black symbols are for instantaneous acclimation; red and blue symbols are for “fast” and “slow” acclimation, respectively, which were modeled using the 10th and 90th percentiles, respectively, of parameters for acclimation kinetics (*τ*_slow_ and *τ*_fast_, the time constants for the slow and fast phases, respectively, of acclimation of *V*_cmax_). Sunfleck length was the same in all cases (2.0 min), whereas shadefleck length was greater at the lower canopy position to reflect the greater fraction of time spent in shadeflecks by lower-canopy leaves.

In very short sunflecks, other factors may dominate dynamics of photosynthesis. For example, buffering of high-frequency (10 – 0.1 Hz) fluctuations in light availability by the ‘capacitance’ afforded by finite metabolite pools can increase the effective light use efficiency of very high sunfleck PPFDs above 100%, as compared to the average photosynthesis rate when the same PPFD is sustained (Pearcy 1990). Rubisco acclimation and deacclimation are probably not relevant to such short sunflecks, which may dominate canopy light regimes under windy conditions, or for plants with very small leaves.

By driving our simulations with observed variation in acclimation kinetics within existing genetic resources for wheat, we were also able to quantify the realistic gains in diurnal carbon capture that should be possible with traditional breeding. We found that the slowest genotypes (modeled using the the highest decile for *τ*_slow_) gained up to 4% less carbon, daily, than the fastest (modeled using the lowest decile for *τ*_slow_), and that the potential gains were greatest for leaves at intermediate canopy positions (LAI = 0.75 and 1.5 m^2^ m^−2^) and for sunflecks of short to intermediate duration (2-16 min) (Fig 6b). Although these potential gains are not as dramatic as the “headline” numbers of 15-20% based on instantaneous acclimation as the target, they are nevertheless worth pursuing and are sufficiently conservative within genotypes to be a feasible target for breeders. We suggest that continuing work should therefore aim to further quantify variation in this important trait across genotypes of wheat, and to identify target genomic regions to assist breeding efforts. In this capacity, we note that our test of the “single-point” *A* vs *c*_i_ method produced very reliable inference of *V*_cmax_ during induction (Fig S6), suggesting that induction kinetics can be reliably quantified with as little as one-tenth the time investment per leaf as required for the full dynamic *A* vs *c*_i_ method used in this study.

## Conclusions

Our study has for the first time identified significant variation in the acclimation time of photosynthesis following dark-to-light transitions across a diverse panel of wheat genotypes, under field and controlled conditions. Slow acclimation of photosynthesis reduced daily carbon assimilation by as much as 16%. These results reinforce the findings of Taylor & Long (2017) in highlighting fast acclimation of photosynthesis, in particular the activation of Rubisco, to fluctuating light as a valuable trait for improvement in wheat breeding programs.

## Supporting Information

### File S1

- Appendix S1. Modeling impact of acclimation kinetics on carbon gain.
- Table S1. List of genotypes.
- Figure S1. Phenological stages at time of measurements.
- Figure S2. Validation of *A*_max_ in TPU-limited conditions.
- Figure S3. Intercellular [CO_2_] at RuBP carboxylation/regeneration transition during induction.
- Figure S4. Relationship between final *A*_max_ and *V*_cmax_ rates and *t*_95_ and *t*_75_ – *t*_25_.
- Figure S5. Effect of leaf orientation on diurnal carbon losses due to slow induction.
- Figure S6. Comparison of default and “one-point” *A* vs. *c*_i_ fitting methods.

### File S2

- R script for dynamic *A* vs *c*_i_ curve fitting
- R script for analysis of acclimation kinetics

### File S3

- CSV file containing extracted kinetics parameters.

## Acknowledgements

This research was supported by the International Wheat Yield Partnership, through a grant provided by the Grains Research and Development Corporation (US00082). TNB was supported by the Australian Research Council (DP150103863 and LP130100183) and the National Science Foundation (Award #1557906). This work was supported by the USDA National Institute of Food and Agriculture, Hatch project 1016439.

## References

Bernacchi CJ, Singsaas EL, Pimentel C, Portis ARJ, Long SP. 2001. Improved temperature response functions for models of Rubisco-limited photosynthesis. Plant, Cell and Environment 24, 253–259.

Busch FA, Sage RF, Farquhar GD. 2018. Plants increase CO_2_ uptake by assimilating nitrogen via the photorespiratory pathway. Nature Plants 4, 46–54.

Carmo-Silva AE, Salvucci ME. 2013. The regulatory properties of Rubisco activase differ among species and affect photosynthetic induction during light transitions. Plant Physiology 161, 1645–1655.

Carmo-Silva E, Scales JC, Madgwick PJ, Parry MAJ. 2015. Optimizing Rubisco and its regulation for greater resource use efficiency. Plant Cell and Environment 38, 1817–1832.

De Kauwe MG, Lin YS, Wright IJ, Medlyn BE, Crous KY, Ellsworth DS, Maire V, Prentice IC, Atkin OK, Rogers A, Niinemets U, Serbin SP, Meir P, Uddling J, Togashi HF, Tarvainen L, Weerasinghe LK, Evans BJ, Ishida FY, Domingues TF. 2016. A test of the ‘one-point method’ for estimating maximum carboxylation capacity from field-measured, light-saturated photosynthesis. New Phytologist 210, 1130–1144.

de Pury DGG, Farquhar GD. 1997. Simple scaling of photosynthesis from leaves to canopies without the errors of big-leaf models. Plant, Cell and Environment 20, 537–557.

Duursma RA. 2015. Plantecophys - An R package for analysing and modelling leaf gas exchange data. Plos One 10, 13.

Farquhar GD, Caemmerer SV, Berry JA. 1980. A biochemical-model of photosynthetic CO_2_ assimilation in leaves of C-3 species. Planta 149, 78–90.

Kromdijk J, Głowacka K, Leonelli L, Gabilly ST, Iwai M, Niyogi KK, Long SP. 2016. Improving photosynthesis and crop productivity by accelerating recovery from photoprotection. Science 354, 857–861.

Lawson T, Vialet-Chabrand S. 2018. Speedy stomata, photosynthesis and plant water use efficiency. New Phytologist.

Long SP, Zhu XG, Naidu SL, Ort DR. 2006. Can improvement in photosynthesis increase crop yields? Plant Cell and Environment 29, 315–330.

Morales A, Kaiser E, Yin X, Harbinson J, Molenaar J, Driever SM, Struik PC. 2018. Dynamic modelling of limitations on improving leaf CO_2_ assimilation under fluctuating irradiance. Plant, cell & environment 41, 589–604.

Mueller-Cajar O, Stotz M, Bracher A. 2014. Maintaining photosynthetic CO_2_ fixation via protein remodelling: the Rubisco activases. Photosynthesis Research 119, 191–201.

Murchie EH, Kefauver S, Ortega JLA, Muller O, Rascher U, Flood PJ, Lawson T. 2018. Photosynthesis: a fast-changing process in an even faster world (Invited Review). Annals of botany 122, 207–220.

Pearcy RW. 1990. Sunflecks and photosynthesis in plant canopies. Annual Review of Plant Physiology and Plant Molecular Biology 41, 421–453.

Portis AR, Salvucci ME, Ogren WL. 1986. Activation of ribulosebisphosphate carboxylase/oxygenase at physiological CO_2_ and ribulosebisphosphate concentrations by Rubisco activase. Plant Physiology 82, 967–971.

Retkute R, Townsend AJ, Murchie EH, Jensen OE, Preston SP. 2018. Three-dimensional plant architecture and sunlit–shaded patterns: a stochastic model of light dynamics in canopies. Annals of botany.

Salter WT, Gilbert ME, Buckley TN. 2018. A multiplexed gas exchange system for increased throughput of photosynthetic capacity measurements. Plant Methods 14, 80.

Seemann JR, Kirschbaum MUF, Sharkey TD, Pearcy RW. 1988. Regulation of ribulose-1,5-bisphosphate carboxylase activity in Alocasia-macrorrhiza in response to step changes in irradiance. Plant Physiology 88, 148–152.

Slattery RA, Walker BJ, Weber AP, Ort DR. 2018. The impacts of fluctuating light on crop performance. Plant Physiology 176, 990–1003.

Soleh MA, Tanaka Y, Kim SY, Huber SC, Sakoda K, Shiraiwa T. 2017. Identification of large variation in the photosynthetic induction response among 37 soybean Glycine max (L.) Merr. genotypes that is not correlated with steady-state photosynthetic capacity. Photosynthesis Research 131, 305–315.

Soleh MA, Tanaka Y, Nomoto Y, Iwahashi Y, Nakashima K, Fukuda Y, Long SP, Shiraiwa T. 2016. Factors underlying genotypic differences in the induction of photosynthesis in soybean Glycine max (L.) Merr. Plant Cell and Environment 39, 685–693.

Taylor SH, Long SP. 2017. Slow induction of photosynthesis on shade to sun transitions in wheat may cost at least 21% of productivity. Philosophical Transactions of the Royal Society B-Biological Sciences 372, 9.

Way DA, Pearcy RW. 2012. Sunflecks in trees and forests: from photosynthetic physiology to global change biology. Tree Physiology 32, 1066–1081.

Woodrow IE, Mott KA. 1989. Rate limitation of non-steady-state photosynthesis by ribulose-1,5-bisphosphate carboxylase in spinach. Australian Journal of Plant Physiology 16, 487–500.

Yamori W, Masumoto C, Fukayama H, Makino A. 2012. Rubisco activase is a key regulator of non-steady-state photosynthesis at any leaf temperature and, to a lesser extent, of steady-state photosynthesis at high temperature. Plant Journal 71, 871–880.

